# In vivo calcium imaging shows that dorsal root ganglion stimulation predominantly activates large-sized sensory neurons in mice

**DOI:** 10.1101/2025.05.27.656338

**Authors:** George L Goodwin, Glenn Franken

## Abstract

Dorsal root ganglion (DRG) stimulation is an effective treatment for patients with refractory neuropathic pain. However, the exact mechanisms through which DRG stimulation exerts its analgesic effects, especially in relation to the recruitment of different soma types within the DRG, remains largely unknown. Here, we examined the effect of DRG stimulation (20Hz) on sensory neuron soma recruitment and neuronal responses induced by mechanical hind paw stimulation using *in vivo* calcium imaging in mice. GCaMP6s was delivered to sensory neurons via an adeno-associated viral vector of serotype 9 (AAV9) injected subcutaneously at P2-5. In adult mice, a stimulation electrode was placed over the L4 DRG, and *in vivo* calcium imaging was performed. The proportion of responsive neurons and response magnitude was assessed during 30-60 seconds of DRG stimulation at different stimulation amplitudes (33%, 50%, 66%, 80%, 100% of motor threshold (MT)) for small, medium and large-sized subpopulations. In a second experiment, the effect of 30 minutes of DRG stimulation was assessed at 66% MT on neuronal responses induced by mechanical hind paw stimulation during and following DRG stimulation. We show that 20Hz DRG stimulation at clinically relevant parameters mainly activates large-size sensory neurons in mice, both in terms of the proportion of neurons responding as well as response magnitude. There was a small decrease in the proportion of neurons responding to mechanical stimulation during DRG stimulation, which did not recover 30 mins after stimulation had been switched off. By contrast, DRG stimulation did not alter the response magnitude of responding neurons.

## Introduction

Dorsal root ganglia (DRG) play a crucial role in the pathophysiology of neuropathic pain (1). The DRG house the cell somata of all three neuronal fiber types (Aβ, Aδ, C), and act as a pivotal relay station for electrical impulses containing information on touch and nociception traveling from the periphery to the spinal cord (2, 3). Following nerve damage, upregulation of neuropeptides, receptors and ion channels can lead to the production of ectopic, aberrant discharges at the level of the DRG (4–9). Such discharges can lead to central sensitization, thereby contributing to the development and maintenance of chronic neuropathic pain (10–12).

DRG stimulation has been used as a last resort treatment approach for patients who do not respond adequately to pharmacological intervention (13, 14). Over the last decade, DRG stimulation has seen consistent use in treating a wide variety of pain disorders (13–17). However, despite frequent use in the clinic, it is still not clear how DRG stimulation exerts its analgesic efficacy.

Initially, it was hypothesized that DRG stimulation might engage gate-control mechanisms in the spinal cord dorsal horn via activation of large sized myelinated Aβ fibers (18–21). In contrast, some studies suggest that DRG stimulation predominantly suppresses the excitability of slow conducting medium sized Aδ or small sized C fibers (22, 23). Given this discrepancy in DRG stimulation literature, more research is needed to investigate which DRG soma types are modulated by DRG stimulation.

*In vivo* calcium imaging is a relatively novel method for studying DRG neuronal function. Using this technique, hundreds of cells can be measured simultaneously (24, 25). In this study, we set out to determine the effect of DRG stimulation at different amplitudes on DRG soma recruitment in GCaMP6s mice. Furthermore, we also assessed the effects of DRG stimulation on neuronal responses induced by mechanical stimulation of the hind paw.

## Methods

### Animals

Thirteen adult C57BL/6J male (n=6) and female (n=7) mice (Charles River, UK) were used in this study. Eight mice were used for *in vivo* imaging and 5 mice for immunohistochemistry. Mice were housed on a 12/12 h light/dark cycle with a maximum of 4 mice per cage, with food and water available *ad libitum*. All experiments were performed in accordance with the United Kingdom Home Office Animals (Scientific Procedures) Act (1986).

### Administration of calcium indicator

We utilized the genetically encoded calcium indicator GCaMP6s for imaging sensory neuron activity (26). GCaMP6s was delivered to sensory neurons via an adeno-associated viral vector of serotype 9 (AAV9), which was administered to mouse pups at P2-P5 as previously described (27). Briefly, groups of 3-4 mice were separated from their mother. 6 µl of AAV9.CAG.GCaMP6s.WPRE.SV40 (Addgene, USA) was injected subcutaneously in the neck, using a 10 µL Hamilton syringe with a 30G needle. Mice were separated from their mother after weaning and then used for *in vivo* imaging or immunohistochemistry from 12 weeks after the injection.

### Immunohistochemistry

Animals were deeply anesthetized with sodium pentobarbital and blood was cleared by transcardial phosphate buffered saline (PBS) perfusion. Lumbar 4 DRGs were dissected out and post-fixed using 4% paraformaldehyde (PFA) for ∼2 hours. Tissue was cryoprotected in 30% Sucrose at 4°C for ≥ 24 hours, embedded in OCT and flash frozen. DRGs were cryosectioned at 9 µm and mounted onto slides. Sections were hydrated in PBS and blocked (6% normal donkey serum, 0.2% TritonX-100, PBS) for 1hr at room temperature (RT). Chicken anti-NeuN (ABN91, Abcam; 1:750) and rabbit anti-GFP (A-11122, Invitrogen; 1:500) antibodies were applied overnight at RT. The next day samples were washed in a wash solution (0.2% TritonX100, PBS) followed by a 2h incubation with Donkey anti-chicken Alexa Fluro 647 and Donkey anti-rabbit Alexa Fluro 488 secondary antibodies (ThermoFisher, 1:1000) at RT. Samples were mounted using Fluoromount-G™ with DAPI (ThermoFisher) and imaged on a confocal microscope (Zeiss LSM 700). Images were analyzed using ImageJ (NIH). Four to 5 section per DRG were analyzed. Neuronal cell bodies were segmented using Cellpose v3.1(28). The cyto2 model was applied with a diameter parameter of 25 pixels to identify regions of interest (ROIs) based on NeuN fluorescence. ROIs that were not circular in morphology were excluded, and any missed neurons were added manually. The neuronal ROI masks generated by Cellpose were exported and imported into ImageJ for fluorescence intensity and soma area measurements. Background fluorescence was estimated from three representative background regions per slide. A neuron was classified as GCaMP6s-positive if fluorescence > background mean + 6 SD. To avoid bias from unequal cell counts, 838 neurons were randomly sampled per animal for histogram analysis.

### *In vivo* calcium imaging

*In vivo* imaging was performed as previously described (25). GCaMP6s was chosen because it is a well-characterized calcium indicator in sensory neurons, with extensive *in vivo* validation by our group and others. Briefly, mice were anesthetized using a combination of drugs: 1-1.25 g/kg 12.5% w/v urethane administered intraperitoneally and 0.5-1.5 % isoflurane delivered via a nose cone. Local anesthetics and non-steroidal anti-inflammatory drugs were not used during preparation of the animal. Body temperature was maintained close to 37°C using a homeothermic heating mat with a rectal probe (FHC). An incision was made in the skin on the back, and the muscle overlying the L3, L4, and L5 DRG was removed. The bone surrounding the L4 DRG was carefully removed in a caudal-rostral direction. Bleeding was prevented using gelfoam (Spongostan™; Ferrosan, Denmark). The exposure was then stabilized at the neighboring vertebrae using spinal clamps (Precision Systems and Instrumentation, VA, USA) attached to a custom-made imaging stage. Finally, the DRG was covered with silicone elastomer (World Precision Instruments, Ltd) to maintain a physiological environment. Prior to imaging, we administered a subcutaneous injection of 0.25 ml 0.9 % sterile saline for hydration. It was then placed under an Eclipse Ni-E FN upright microscope (Nikon). The ambient temperature during imaging was kept at 32°C throughout. All images were acquired using a 10× dry objective. A 488-nm Argon ion laser line was used to excite GCaMP6s, and the signal was collected at 500–550 nm. Time lapse recordings were taken with an in-plane resolution of 512 × 512 pixels and a partially (∼2/3rd) open pinhole for confocal image acquisition. The optical section thickness for these recordings was estimated to be between 80-100µm. All recordings were acquired at 3.65 Hz.

### DRG stimulation

Bipolar electrodes were carefully placed either side (caudal and rostral) of the left L4 DRG using a microscope on the dissection table (See Figure 1A for a diagram of where electrodes were typically positioned). The DRG were stimulated with a constant current stimulator (Digitimer, Welwyn Garden City, United Kingdom), coupled with a voltage stimulator (Pulsemaster A300, World Precision Instruments, Ltd). Following electrode placement, the motor threshold (MT) was accurately determined by gradually increasing the stimulation intensity using a frequency of 2 Hz and a pulse width of 200 µs. The MT was defined as the current inducing observable contractions of the ipsilateral lower trunk or hind limb. The mouse was then placed onto the microscope stage and the MT was re-checked to ensure that the electrodes had not changed position whilst moving the mouse. 20 Hz DRG stimulation with a 200 µs pulse width was applied for 30s-60s at 33%, 50%, 66%, 80%, 100% MT, with 1-minute in between each stimulation for recovery in n=5 experiments (29). In two experiments, only 33-80% MT was tested (Figure 1B). The order of stimulation was randomized for each experiment using randomize.org. For the mechanical stimulation studies, DRG stimulation was delivered for 30 minutes at 20 Hz, using a pulse width of 200 µs and an amplitude of 66% MT. These parameters are reflective of clinical DRG stimulation, where frequencies of 20 Hz, pulse widths between 200-300 µs, and stimulation amplitudes just above the sensory threshold are typically used (29).

**Figure 1.**
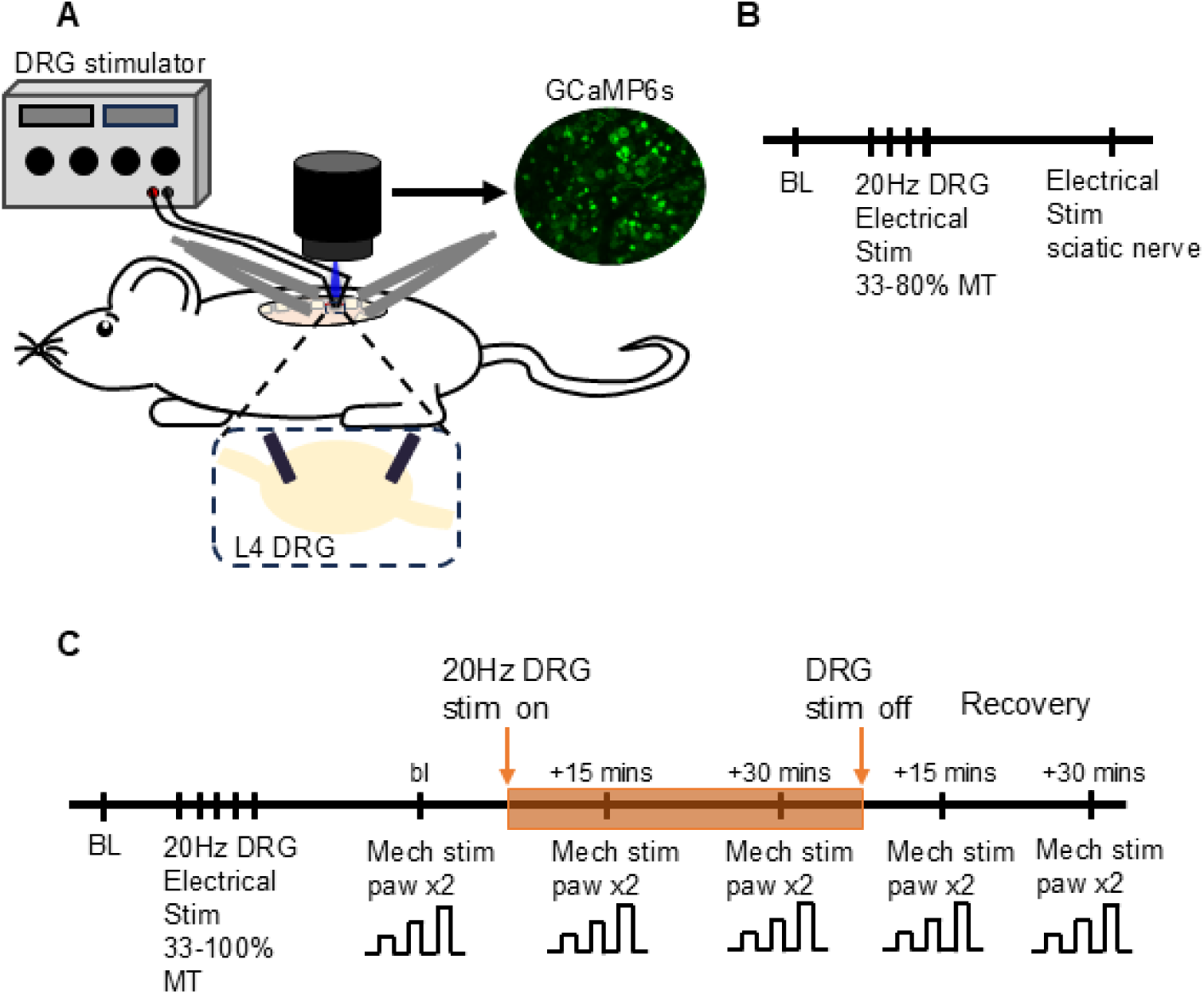
Diagram of experimental set up and stimulation paradigms. A) Schematic showing experimental set up. A constant current stimulator was used to deliver a pulse of 20Hz stimulation to the L4 Dorsal root ganglion (DRG). Responses were recorded from neurons labelled with GCaMP6s using a confocal microscope. B) Schematic of experimental paradigm 1, n=2 mice. C) Schematic of experimental paradigm 2, n=5 mice.

### Mechanical stimulation of sensory afferents

In some mice (n=5), A custom-made mechanical stimulator was used to mechanically stimulate sensory afferents in the hind paw (See (30) for details). The paw was positioned face up and was fixed into position using blue tack. The tip of the mechanical stimulator, which contains 6×4 blunt pins (2mm in height and measuring approximately 6×4mm in total), was positioned 1-2mm above the medial plantar surface of the hind paw, an area to which many L4 afferents project in mice (31). Three forces of 200g (1.96N, 0.08N/pin), 500g (4.90N, 0.20N/pin) and 800g (7.85N, 0.33N/pin) were applied sequentially for 1s duration, allowing 1 minute in between each stimulation for recovery. Each series of force steps was repeated twice at baseline and at different timepoint during and after stimulation of the DRG (Figure 1C). To optimize neuron detection by the activity algorithm in Suite2p, we applied a pinch stimulus to the hind paw using serrated forceps, maximizing the number of neurons automatically identified.

### Imaging data analysis

Timelapse recordings were scaled to 8-bit in Fiji/ImageJ, Version 1.53. The image analysis pipeline Suite2P (v 0.9.2) (32) was utilized for motion correction, automatic region of interest (ROI) detection and signal extraction. ROIs that were not automatically detected in suite2p were added manually. Further analysis was undertaken with a combination of Microsoft Office Excel 2013, and Python (Version 3.9). A region of background was selected, and its signal subtracted from each ROI. To generate normalized data, the following calculation was used: ΔF/F0 = (Ft-F0)/F0, where Ft is the fluorescence at time t and F0 is the fluorescence average over the first 30 seconds of the recording. ΔF/F0 is expressed as a ratio. To correct for fluctuations in F0 values over time, such as minor drifts in the fluorescent baseline, ΔF/F0 values for each event were recalculated using F0 values averaged from the 10 frames immediately preceding the event onset. To avoid aberrant amplification because of small F0 values, cells with very low basal fluorescence after background subtraction (<1.5 A.U. from 0-255 on an 8-bit scale) were excluded. Two hundred and seventy two out of 1725 neurons were excluded (average = 15.8% /experiment).

The area of each neuron in the L4 DRG was estimated by using the radius output from Suite2p. Cells were grouped into small (<600µm^2^), medium (≥600 - <1000µm^2^) and large-sized (≥1000µm^2^) neurons. Note that cell areas measured using immunohistochemistry will appear smaller than those measured using *in vivo* imaging, due to tissue shrinkage during cryopreservation and because histological sections capture only a slice of each neuron rather than the full soma. Cells that could not clearly be identified and had substantial overlap with neighboring cells were excluded from the analysis.

Spontaneous activity in neurons i.e. that which is generated without any apparent cause, was assessed using a trained machine learning classifier during the first 2.5 minutes of the recording (25). Only experiments with at least 2.5 minutes of baseline were included (n = 5 mice).

For electrical stimulation, a response was considered positive if the mean signal during stimulation was 8*SD ≥ the relative baseline. For mechanical stimulation, a response was positive if the maximum signal was 6*SD ≥ the relative baseline. For each size subgroup, the percentage of neurons responding to a stimulus was calculated by dividing the number of responders by the total number of neurons in that size subgroup. For responding neurons, the average magnitude of the response during the stimulation period was calculated.

There were a small proportion of neurons that responded to mechanical stimulation and 66% MT DRG stimulation in experiments examining the changes to mechanically induced responses. These neurons were excluded from the mechanical response analysis because the responses during DRG stimulation could not be accurately measured I.e The ΔF/F0 for these neurons was reduced because the baseline level of fluorescence was raised by DRG stimulation. Sixty-nine out of 1000 neurons were excluded (average = 6.5%/experiment).

### Data loss and exclusion

One experiment was excluded because the GCaMP6s labelling was very poor. The data from this experiment was not analyzed.

### Quantification and statistical analysis

Graphing and statistical analysis was undertaken with a combination of Excel 2013, Python (Version 3.9), GraphPad Prism (Version 10) and SPSS (Version 29.0). We used a linear mixed-effects model (LMM) to analyze the effects of stimulation strength and neuron size on responses and to assess effects of DRG stimulation on mechanically induced responses. Details of sample sizes are recorded in the appropriate figure legend.

## Results

The genetically encoded calcium indicator GCaMP6s was delivered to sensory neurons via a viral approach. Immunohistochemical labelling was performed to verify that the virus transduced all sizes of sensory neurons within the DRG used for *in vivo* imaging. The soma of all neuronal size classes exhibited GCaMP6s expression (Figure 2 A&B). However, there was a slight bias towards small to medium sized neurons: on average, 60.4 +/- 2.7% of neurons with soma areas <700 µm² expressed GCaMP6s, compared with 35.5 +/- 4.1% of neurons ≥700 µm (Figure 2C).

**Figure 2.**
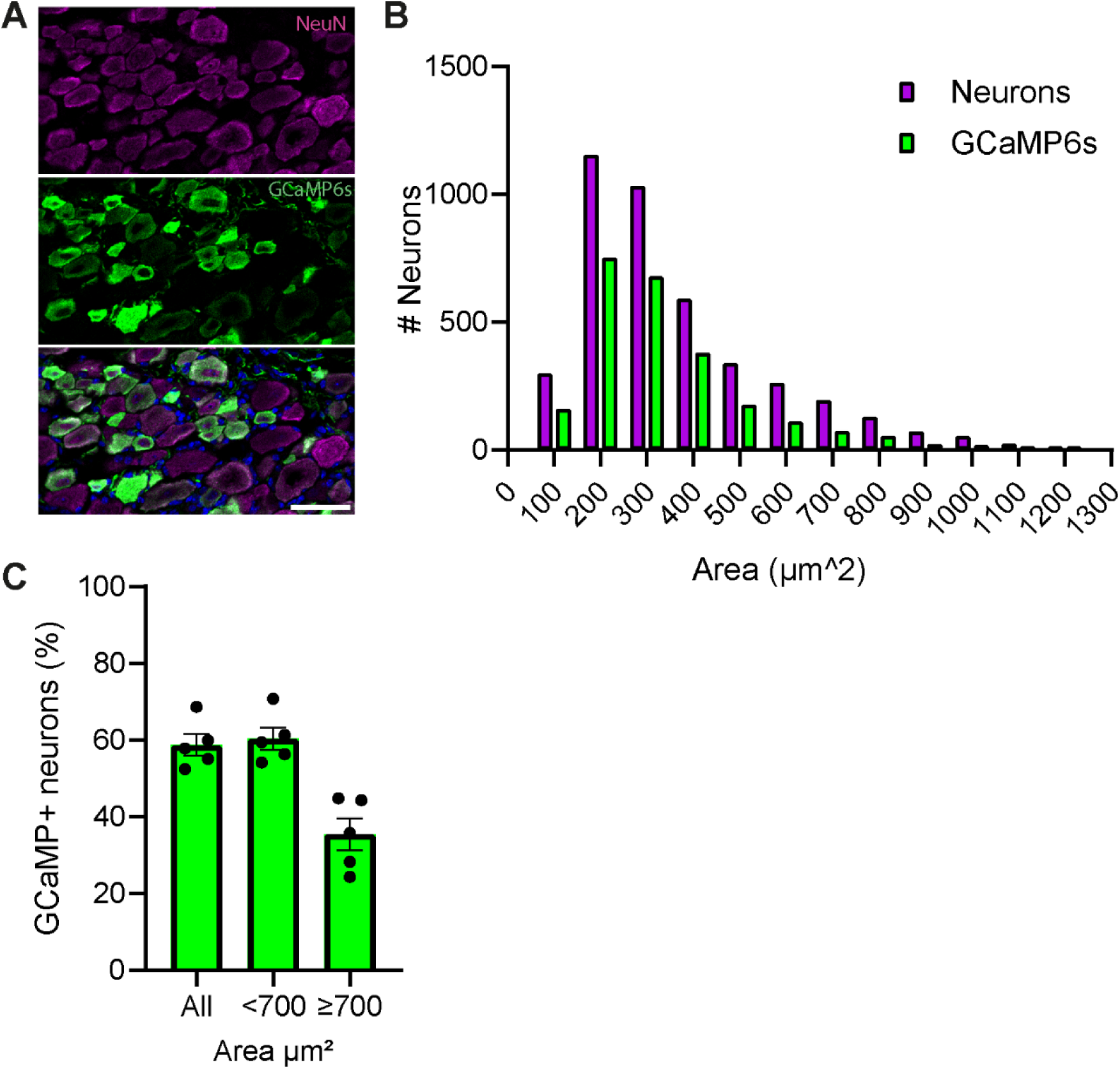
AAV9-GCaMP6s delivery to mouse pups labels sensory neurons of all sizes classes. A) Example immunolabelled section of the L4 DRG from a mouse injected with AAV9-GCaMP6s at p3. An anti-NeuN antibody was used to label sensory neurons (magenta - top panel) and an anti-GFP antibody was used to label GCaMP6s labelled sensory neurons (green - middle panel). The bottom panel shows a merge of the two channels with DAPI staining (blue). Scale bar = 50 µm. B) Size histogram showing the number of neurons (magenta) and GCaMP labelled neurons (green) in bin widths of 100µm². The label denotes the upper bin limit. N=4190 neurons from 5 L4 DRGs. C) Graph shows the percentage of neurons that were GCaMP6s +. N=5 L4 DRGs from 5 Mice; Mean +/- SEM.

A total of 1453 neurons were recorded from n=7 mice in *in vivo* imaging experiments. Neuronal soma sizes measured from *in vivo* recordings were larger compared to that assessed via immunohistochemistry (Supplementary Figure 1). This is likely due to tissue shrinkage during cryopreservation and because histological sections capture only a slice of each neuron rather than the full soma. Based on the size distribution (Supplementary Figure 1), we defined boundaries at 600 µm² and 1000 µm² to classify neurons into small–medium and medium–large size categories for analysis of *in vivo* imaging experiments.

For DRGS, electrodes were carefully positioned on the surface of the L4 DRG. A low number of neurons exhibited spontaneous activity during the first 2.5 minutes of the recording (mean 12.9%, +/- 1.6%, n=5 mice), suggesting that electrode placement did not induce inflammation-related neuronal sensitization. The mean motor threshold was 50.6 µA (+/- 9.1, n=7 mice; Figure 3A). Following threshold determination, GCaMP6s-labelled DRG neurons were imaged using a confocal microscope. Electrical stimulation was applied at varying intensities relative to the MT to identify which types of DRG neurons responded. As expected, increasing the stimulation strength caused an increase in the proportion of responding neurons (F (4,72) [Stimulation strength] = 45.5, p<0.0001, linear mixed model; Figure 3 B-D). At every stimulation strength tested, large sized neurons were preferentially activated in comparison to small and medium sized neurons (F (2,72) [Size] = 107, p<0.0001, linear mixed model, n=7). At 66% of the MT, a commonly used stimulation intensity in preclinical pain behavior experiments that resembles clinical settings, 31.1% (± 6.8, n=7) of large-sized neurons were activated (Figure 3 B-D). In contrast, fewer small- and medium-sized neurons were activated at 66% MT (6.5% +/- 1.9 small neurons; 16.2% +/- 5.2 medium neurons, n=7). Similarly, the response magnitudes during DRG stimulation were greater in large sized neurons compared to small and medium sized neurons (F (2,60) [Size] = 31.3, p<0.0001, linear mixed model, n=7; Supplementary Figure 2). Electrical stimulation at 66% of the MT increased the GCaMP fluorescence of large-sized neurons by 3.7 fold (+/- 0.47, n=7), while small increases occurred in small and medium-sized neurons (1.3 fold +/-0.2 and 2.9 fold +/-0.5, respectively, Supplementary Figure 2).

**Figure 3.**
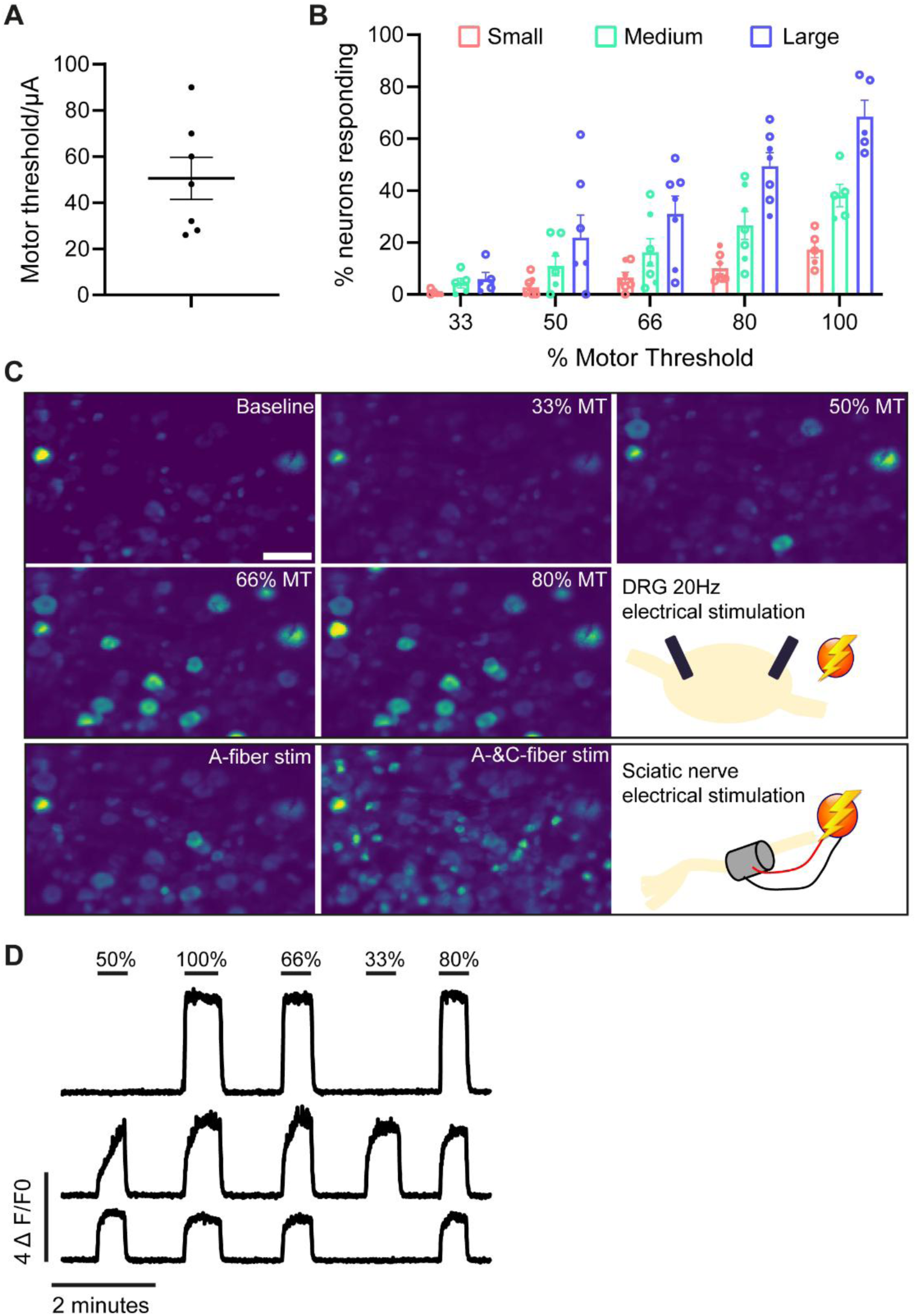
20Hz DRG stimulation mostly activates large sized neurons when stimulating between 33%-100% of the motor threshold. A) Graph showing the motor threshold for each experiment. Each dot represents an experiment. B) The proportion of neurons in each size subcategory responding to 33-100% of motor threshold (MT) 20hz DRG stimulation. n=5 mice for 33-100% MT and n=7 mice for 33-80% MT stimulation. N=2 males (closed circles) and n=5 females (open circles). n=123–274 neurons/mouse. C) Example average intensity projections taken from 30s clips during different experimental periods. Top panels show average fluorescence intensities at baseline and at different DRG stimulation strengths. Bottom panels show average fluorescence intensities in response to stimulation of the sciatic nerve at low threshold (left) and suprathreshold (right) stimulation strengths. Scale bar = 100µm. D. Three example recordings showing the calcium responses of large-sized neurons responding to DRG stimulation at different stimulation amplitudes.

DRG stimulation at 20Hz is reported to induce analgesic responses in both rodents and humans (13–17, 33–35). However, the mechanisms of analgesia are not completely understood. To determine if DRG stimulation reduces neuronal activity within cells bodies of the DRG, mechanically induced responses were measured before, during and after 20Hz DRG stimulation. Neurons in the hind paw ipsilateral to the DRG being imaged were stimulated using a mechanical stimulator (Figure 1C). As expected, increasing force activated a greater number of neurons (F(2,176) [Force] = 114, p < 0.0001, Linear mixed model, n = 5) and increased their response magnitude (F(2,167) = 44, p < 0.0001, Linear mixed model, n = 5; Figure 4A-D). Following application of DRG stimulation, the graded nature of responses to increasing force remained consistent over time F(8,167) [Force*Time] = 0.39, p =0.93, Linear mixed model, n = 5). Similarly, there was no significant change in the response magnitude of the responding neurons following 20Hz DRG stimulation (F (4,167) [Time] = 0.2, p=0.93, linear mixed model; n=5; Figure 4D). By contrast, the proportion of neurons responding to mechanical stimulation significantly decreased following DRG stimulation (F(4,176) [Time] = 2.5, p = 0.042, linear mixed model, n = 5; Figure 4C). The proportion of neurons responding to mechanical stimulation declined after DRG stimulation (Mean difference: Baseline vs. +30 min = 5.2, SE = 2.3, 95% CI: -1.4 to 11.7), but responsivity did not recover after stimulation ceased (Mean difference: +30 min vs. Recovery +30 min = 0.5, SE = 2.3, 95% CI: -6 to 7). Pairwise comparisons with Bonferroni correction did not reveal significant differences between individual time points (all p>0.05). The reduction in the proportion of responders was consistent across all neuron size groups and forces, as there were no significant interactions between time and neuron size (F(8,176) = 0.16, p = 0.99) or time and force (F(8,176) = 1.2, p = 0.32; linear mixed model).

**Figure 4.**
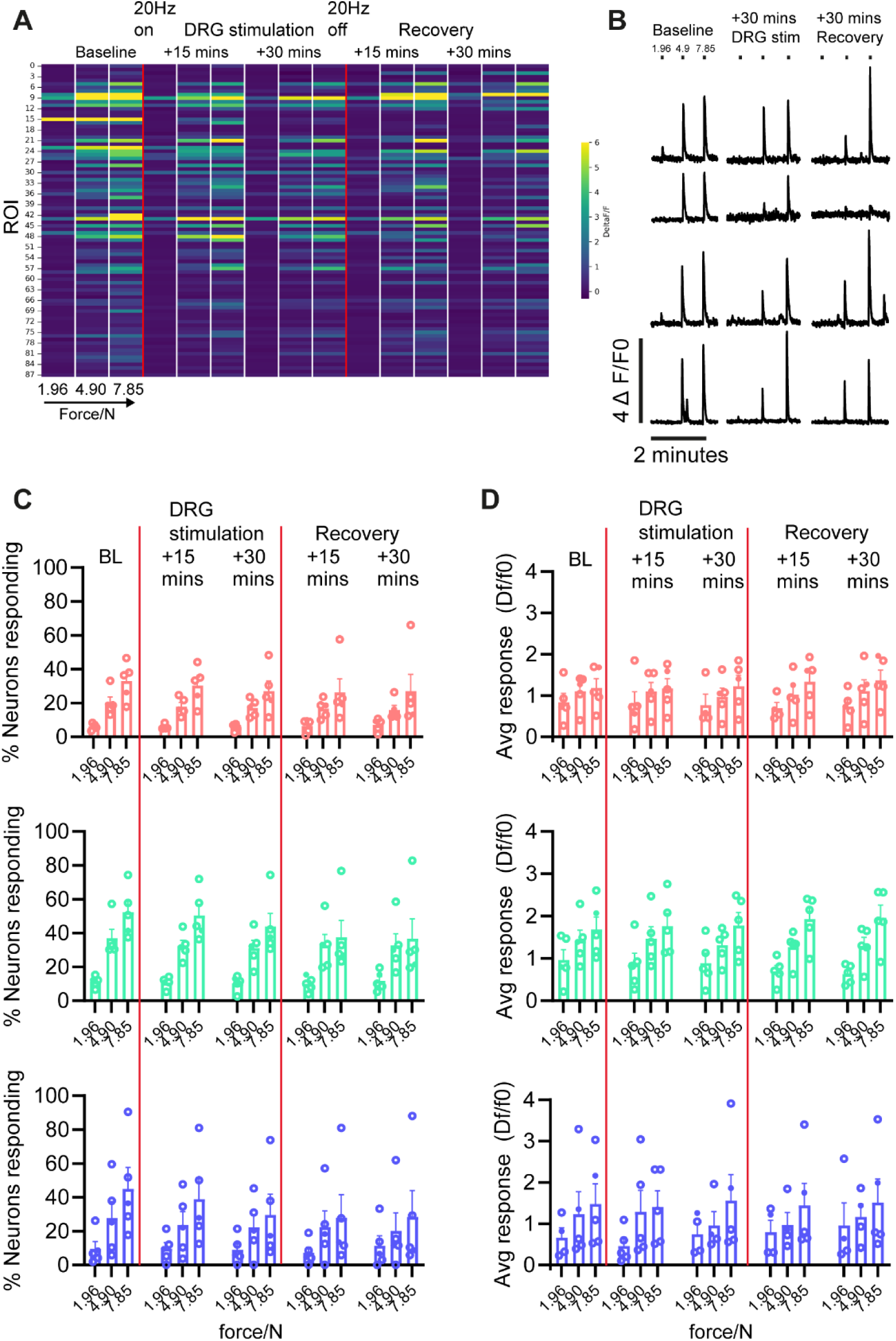
20Hz DRG stimulation did not attenuate mechanically induced responses generated in hind paw. A) Heat map showing the DeltaF/F0 normalized data for an example experiment. Each row shows data from a neuron that responded to mechanical stimulation of the paw during the baseline period. Each column represents the average DeltaF/F0 during mechanical stimulation of the hind paw. B) Four example recordings showing the calcium responses of neurons responding to mechanical stimulation before, during and after DRG stimulation. C) and D) Graphs show the proportion of neurons responding (C) and average responses (D) to mechanical stimulation of the hind paw pre, during and post 20Hz DRG stimulation. Top panel = small sized neurons, middle panel = medium sized neurons, Bottom panels = large sized neurons. Mean +/- SEM.

## Discussion

In this study, we assessed the effect of DRG stimulation at different stimulation amplitudes on DRG soma recruitment. We found that DRG stimulation at clinically relevant amplitudes mainly recruits large sized neurons, and to a lesser extent medium sized neurons. In contrast, only very few small-sized neurons were recruited. The proportion of neurons responding to mechanical stimulation decreased following DRG stimulation, but this responsivity did not recover after stimulation ceased.

Using *in vivo* calcium imaging we found that DRG stimulation mainly recruits large sized A-somata across all stimulation amplitudes. At clinically relevant amplitudes of 66% MT, 31.1% of large-sized neurons were activated, whereas relatively few small- (6.5%) and medium-sized (16.2%) neurons were activated. Consistent with this finding, Chao *et al.* reported a negative correlation between cell size and DRG stimulation intensity needed to activate neuronal cell bodies(23). However, this study found no relationship between DRG stimulation evoked response magnitude and cell diameter *in vivo*, contrasting our findings: at 66% MT, large-sized neurons showed a 3.7 fold increase in response magnitude, compared to 2.9-fold in medium and 1.3-fold in small neurons. This discrepancy might be due to the difference in recording set ups: we employed a ‘dry’ set up, whereas the DRG was perfused in imaging experiments conducted by Chao *et al.,* meaning that the current might have dissipated into the surrounding solution. Our findings are consistent with a computational study showing that DRG stimulation drives the activity of large myelinated Aβ-fibers, but not C-fibers (17). The importance of large-diameter fiber activation was further underscored by Ting *et al.*, who demonstrated that DRG stimulation evokes activity in Aα/β-fibers of the sciatic nerve (19). Interestingly, a recent study showed that the density of small, myelinated axons is lowest in the dorsal region of the middle of the DRG. Hence, stimulation near the middle of the ganglion may modulate large axons, while minimizing activation of small myelinated nociceptive fibers (36). Collectively, these data suggest that DRG stimulation preferentially activates large-diameter fibers, implicating their importance in DRG stimulation mediated analgesia.

Our mechanical stimulation experiments showed that DRG stimulation does not affect the response magnitude – an indicator of the level of neuronal excitability - in responding neurons. These results appear to contrast a previous study reporting that DRG stimulation reduces cell soma excitability, as assessed by examining conduction velocity (37). However, this could not be assessed in our study using GCaMP6s due to its relatively slow kinetics (26). Examining kinetics would have required a faster GCaMP variant, such as GCaMP8f, but this was not feasible due to limitations in our microscope’s imaging speed. By contrast, the proportion of neurons responding to mechanical stimulation decreased following DRG stimulation. It is important to note that this responsivity did not recover after stimulation ceased. As the reduction in the proportion of responders was consistent across all neuron size groups and forces, it may reflect repeated mechanical stimulation inducing inflammation (38), which has been reported to reduce neuronal responsivity (39). Although we did not observe a substantial reduction of the number of neurons responding to mechanical stimulation at the level of the cell soma, other studies, which examine transmission at the level of the dorsal root, have reported conduction block (23, 40, 41). Specifically, transmission was blocked by DRG stimulation in C-fibers, and to a lesser extent Aδ and Aβ-fibers (23). This data is further supported by computational work that shows that DRG stimulation can amplify the natural filtering effect of the DRG, thereby blocking afferent signaling and ectopic activity (22).

DRG stimulation was originally developed as an alternative to spinal cord stimulation (SCS). SCS is known to activate non-nociceptive Aβ fibers in the spinal cord dorsal columns and engage gate-control mechanisms in the spinal cord dorsal horn, thereby inhibiting incoming transmission from nociceptive C-fibers (42). In theory, DRG stimulation might engage similar gate-control mechanisms, through direct activation of Aβ somata in the DRG. Indeed, our study shows that large-sized somata, which encompass Aβ somata, are predominantly activated by DRG stimulation. Another theory is that local T-junction filtering occurs at the level of the DRG. DRG somata have a unique pseudo-unipolar design, with one common axon (stem) that branches into two separate axons, connected by T-junction that connects the branching point to the cell body (1). DRG stimulation is proposed to depolarize the somatic membrane, thereby creating a reduction in membrane excitability at the T-junction (23, 37). This loss in membrane excitability at the site of the T-junction is thought to be Ca^2+^ dependent and involve Ca^2+^ -sensitive K ^+^ channels, producing hyperpolarization in the axon stem and T-junction (22, 23, 43, 44). Regardless of the precise soma type that is modulated by DRG stimulation, it is reasonable to expect that ultimately, this leads to a reduction in output of second order projection neurons and pain transmission to supraspinal centers. Indeed, a previous studies showed that DRG stimulation can reduce spontaneous activity in wide-dynamic-range neurons (45) and attenuate blood oxygen-level dependent (BOLD) responses in areas related to sensory and pain-related responses (46).

Our study has limitations. The mechanical stimulation experiments were performed in 5 mice. Given this sample size and the variability observed, we were only able to detect very large effect sizes. In contrast, previous behavioral studies typically used 8-12 rats to detect an analgesic effect (21, 33–35, 47–50). We did not include a sham DRG stimulation control group and therefore we were not able to determine if the progressive decline in mechanical responsivity was an effect mediated by DRG stimulation or repeated mechanical testing. It is important to note that we were only measuring excitability at the level of the soma and so it is entirely possible that the T-junction filtering mechanisms described above, may have caused inhibition of neurons along the dorsal root, which we would have missed. We delivered GCaMP6s to sensory neurons via a viral approach and used this as a proxy of neuronal excitability. Viral delivery via AAV9 labels most neuronal subtypes, including Aα and Aβ fibers (51), but it is important to note that relatively few tyrosine hydroxylase (TH) positive neurons, which are markers of low threshold mechanoreceptor C-fibers (C-LTMR), are labelled using this approach (52). Therefore, it is likely that the C-somata activated in our preparations were likely nociceptive and it is not known whether C-LTMRs would be activated. Of note, activation of this subclass of C-fibers and subsequent engagement of the endogenous opioid system was recently suggested to underly DRG stimulation induced analgesia, especially at low frequencies (<20Hz)(53, 54).

In conclusion, our study showed that DRG stimulation at clinically relevant amplitudes mainly recruits large sized A-somata, and to a lesser extent medium sized A-somata and C-somata. The proportion of neurons responding to mechanical stimulation significantly decreased following DRG stimulation, but this responsivity did not recover after stimulation ceased. These results imply that both DRG stimulation induced spinal gating mechanisms as well as local filtering mechanisms should be considered. Future studies should further investigate the effect of DRG stimulation on specific subtypes of DRG somata, including LTMR C-somata and a range of DRG stimulation parameters.

## Declarations

### Ethics approval

All experiments were performed in accordance with the United Kingdom Home Office Animals (Scientific Procedures) Act (1986).

### Availability of data and materials

Raw data and code for doing calcium imaging analysis will be shared in the OSF under accession number:

### Authors’ contributions

Author contributions: G.G. conceived, designed, and performed experiments, analyzed data, and wrote the manuscript. G.F conceived, designed, and performed experiments, analyzed data, and wrote the manuscript.

## Acknowledgements

We would like to thank Dr. Martyn Jones for use of his experimental equipment. We thank Dr. Kim Chisholm for the illustration of the mouse used in Figure 1.

## Authors’ information

This research was funded in whole or in part by UK Research & Innovation. For the purpose of Open Access, the author has applied a CC BY public copyright license to any Author Accepted Manuscript (AAM) version arising from this submission.

## Figures

**Supplementary figure 1.**
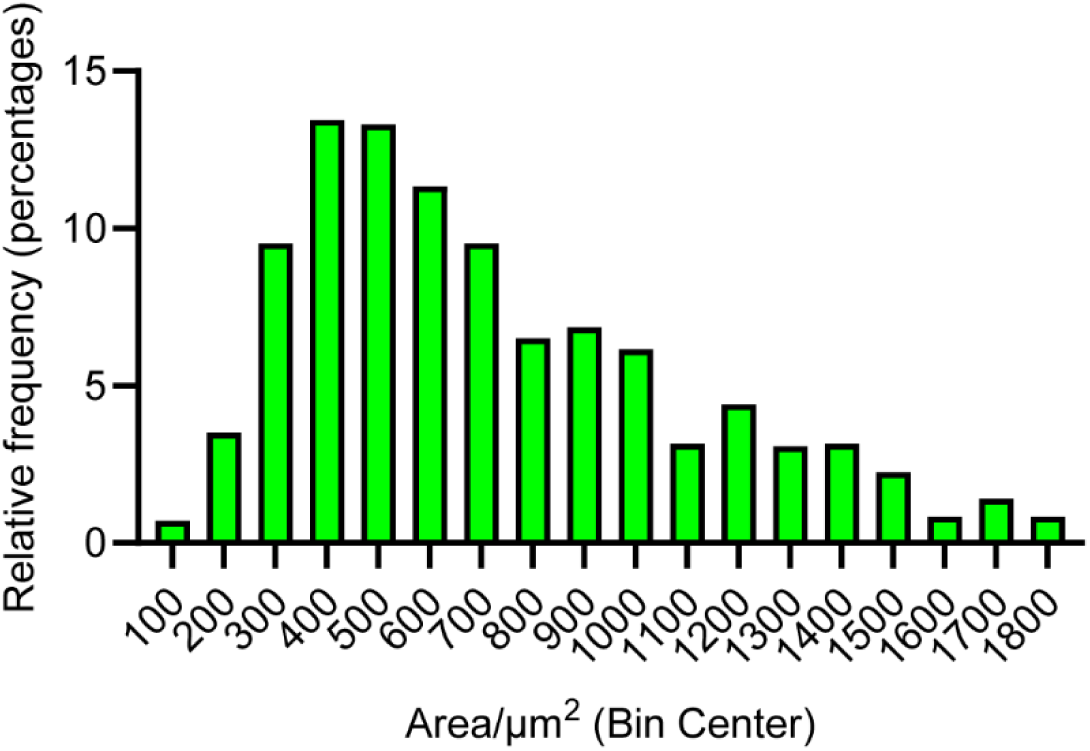
Soma size distribution of neurons imaged *in vivo*. The histogram depicts the distribution of neuronal soma areas obtained from *in vivo* calcium imaging experiments. n= 1453 neurons from n=7 mice.

**Supplementary figure 2.**
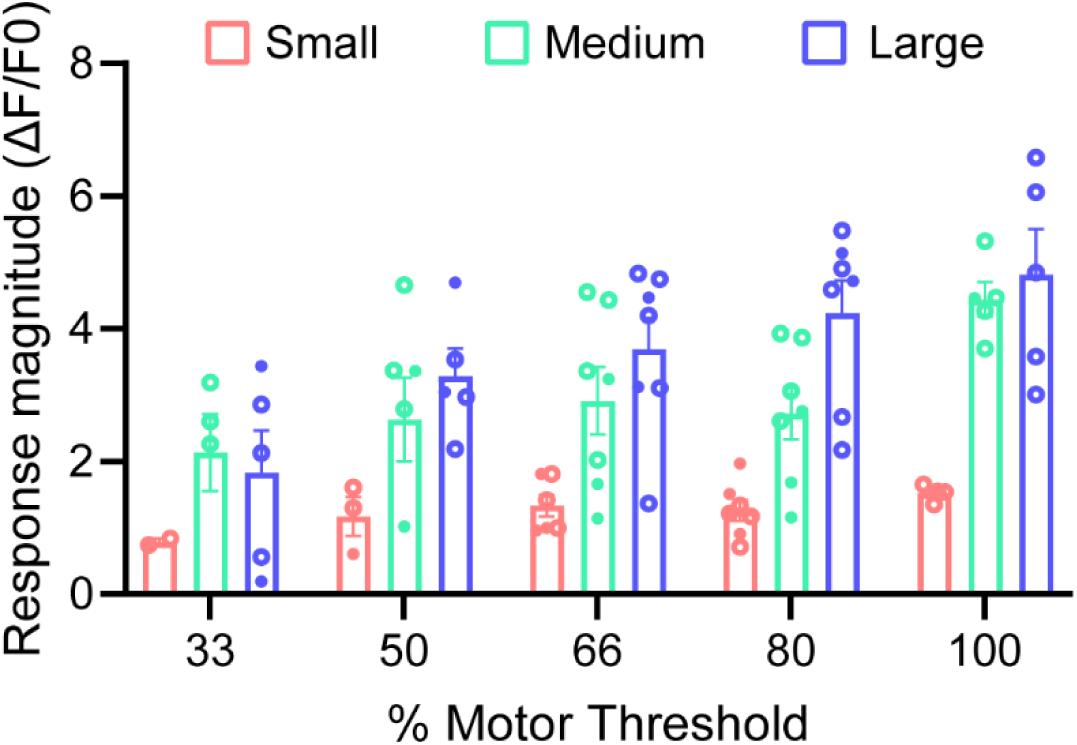
Response magnitudes following 20Hz DRG stimulation between 33%-100% of the motor threshold. Graph shows the average magnitude of response during DRG stimulation for small, medium and large sized neurons. Mean +/- SEM.

